# Computational Causal Modeling of the Dynamic Biomarker Cascade in Alzheimer’s Disease

**DOI:** 10.1101/313353

**Authors:** Jeffrey R. Petrella, Wenrui Hao, Adithi Rao, P. Murali Doraiswamy, for the Alzheimer’s Disease Computational Modeling Initiative

**Author notes:** Send Correspondence to Jeffrey R. Petrella, MD, Department of Radiology, DUMC - Box 3808, Durham, NC 27710-3808.

## Abstract

**Background:** Alzheimer’s disease (AD) is a major public health concern and there is an urgent need to better understand its complex biology and develop effective therapies. AD progression can be tracked in patients though validated imaging and spinal fluid biomarkers of pathology and neuronal loss. We still, however, lack a coherent quantitative model that explains how these biomarkers interact and evolve over time. Such a model could potentially help identify the major drivers of disease in individual patients and simulate response to therapy prior to entry in clinical trials. A current theory of AD biomarker progression, known as the dynamic biomarker cascade model, hypothesizes AD biomarkers evolve in a sequential, but temporally overlapping manner. A computational model incorporating assumptions about the underlying biology of this theory and its variations would be useful to test and refine its accuracy with longitudinal biomarker data from clinical trials.

**Methods:** We implemented a causal model to simulate time-dependent biomarker data under the descriptive assumptions of the dynamic biomarker cascade theory. We modeled pathologic biomarkers (beta-amyloid and tau), neuronal loss biomarkers and cognitive impairment as non-linear first order ordinary differential equations (ODEs) to include amyloid-dependent and non-dependent neurodegenerative cascades. We tested the feasibility of the model by adjusting its parameters to simulate three specific natural history scenarios in early-onset autosomal dominant AD and late-onset AD, and determine whether computed biomarker trajectories agreed with current assumptions of AD biomarker progression. We also simulated the effects of anti-amyloid therapy in late-onset AD.

**Results:** The computational model of early-onset AD demonstrated the initial appearance of amyloid, followed by biomarkers of tau and neurodegeneration, followed by onset of cognitive decline based on cognitive reserve, as predicted by prior literature. Similarly, the late-onset AD computational models demonstrated the first appearance of amyloid or non-amyloid-related tauopathy, depending on the magnitude of comorbid pathology, and also closely matched the biomarker cascades predicted by prior literature. Forward simulation of anti-amyloid therapy in symptomatic late-onset AD failed to demonstrate any slowing in progression of cognitive decline, consistent with prior failed clinical trials in symptomatic patients.

**Conclusion:** We have developed and computationally implemented a mathematical causal model of the dynamic biomarker cascade theory in AD. We demonstrate the feasibility of this model by simulating biomarker evolution and cognitive decline in early and late-onset natural history scenarios, as well as in a treatment scenario targeted at core AD pathology. Models resulting from this causal approach can be further developed and refined using patient data from longitudinal biomarker studies, and may in the future play a key role in personalizing approaches to treatment.

## Introduction

Alzheimer’s disease (AD), one of the leading public health priorities in the U.S., is projected to affect over 15 million people by 2050. The high failure rate of clinical drug trials over the past decade is in large part rooted in an incomplete understanding of its complex causal mechanisms (3). Genetic pathway analyses implicate over 1000 different molecular species and over 30 metabolic pathways in the pathophysiology of AD, including amyloid and tau proteinopathies, inflammation, microglial activation, alterations in signaling pathways, cholesterol metabolism, and cholinergic function (4). It is therefore likely that AD is not a single disease, but a common end-stage pathway resulting from multiple interacting etiologies. Effective treatment will likely require a personalized medicine approach to track disease progression, determine the major pathophysiologic drivers and tailor an appropriate therapy.

AD progression can be tracked in patients, from pre-symptomatic to late-stage disease, through several validated biomarkers. These are biomarkers of AD core pathology (cerebrospinal fluid and PET scan markers of beta-amyloid and tau proteins), and biomarkers of neuronal loss (FDG-PET and volumetric MR imaging). Data from the Alzheimer’s Disease Neuroimaging Initiative (ADNI), and other naturalistic studies, have led to a hypothetical model of disease progression known as the dynamic AD biomarker cascade theory (5), which hypothesizes that AD biomarkers evolve in a sequential, but temporally overlapping manner. According to the hypothesis, amyloid pathology is an early event, leading to tau pathology, followed by neuronal loss and cognitive decline. Additional refinements of the model have been proposed to make it more generalizable to community-based aging populations, including the addition of suspected non-amyloid pathology (SNAP) (e.g., cerebrovascular disease, age-related changes and non-AD tauopathies), as well as the concept of cognitive reserve (e.g. protective factors such as genetics or education), both of which could influence the variability in onset of biomarkers and cognitive decline (2). Although this theoretical model has been operationalized into a categorical scheme for classifying patients, the system remains descriptive, and makes no assumptions regarding putative causal relationships among biomarkers (6). Understanding how these biomarkers interact, evolve over time and result in cognitive expression of disease will be essential to harness them in a personalized medicine approach to AD diagnosis and treatment. Given the complexity of AD, a rigorous mathematical and computational modeling approach, such as that offered by systems biology, will be a critical component.

The tools of systems biology may be used to incorporate clinical biomarkers of disease progression into a computational model to determine the major patho-etiologic drivers of disease in individual patients and help simulate the effects of potential interventions. One modeling approach, known as causal modeling, refers to an explicitly formulated mathematical description of the biological phenomena of interest, based on existing knowledge, in terms of cause and effect relationships (7). This is in contrast to correlative models which merely describe statistical associations between variables without regard to the mechanism driving the phenomena under investigation. Our goal was to construct and test the feasibility of an initial computational causal model (CCM) of AD biomarker progression, based on the updated dynamic AD biomarker cascade theory (1, 2). This would enable the theory to be tested rigorously with existing data and further refined as new data becomes available.

## Methods

For the construction of the causal model, assumptions about biomarker relationships and temporal course were drawn from prior literature (1, 2, 5). We tested whether the CCM would lead to the predicted biomarker trajectories described in the literature, and whether it would predict failed outcome of anti-amyloid therapy started late in the disease course (8). Figure 1 shows the variables and their relationships in the computational model.

**Figure 1:**
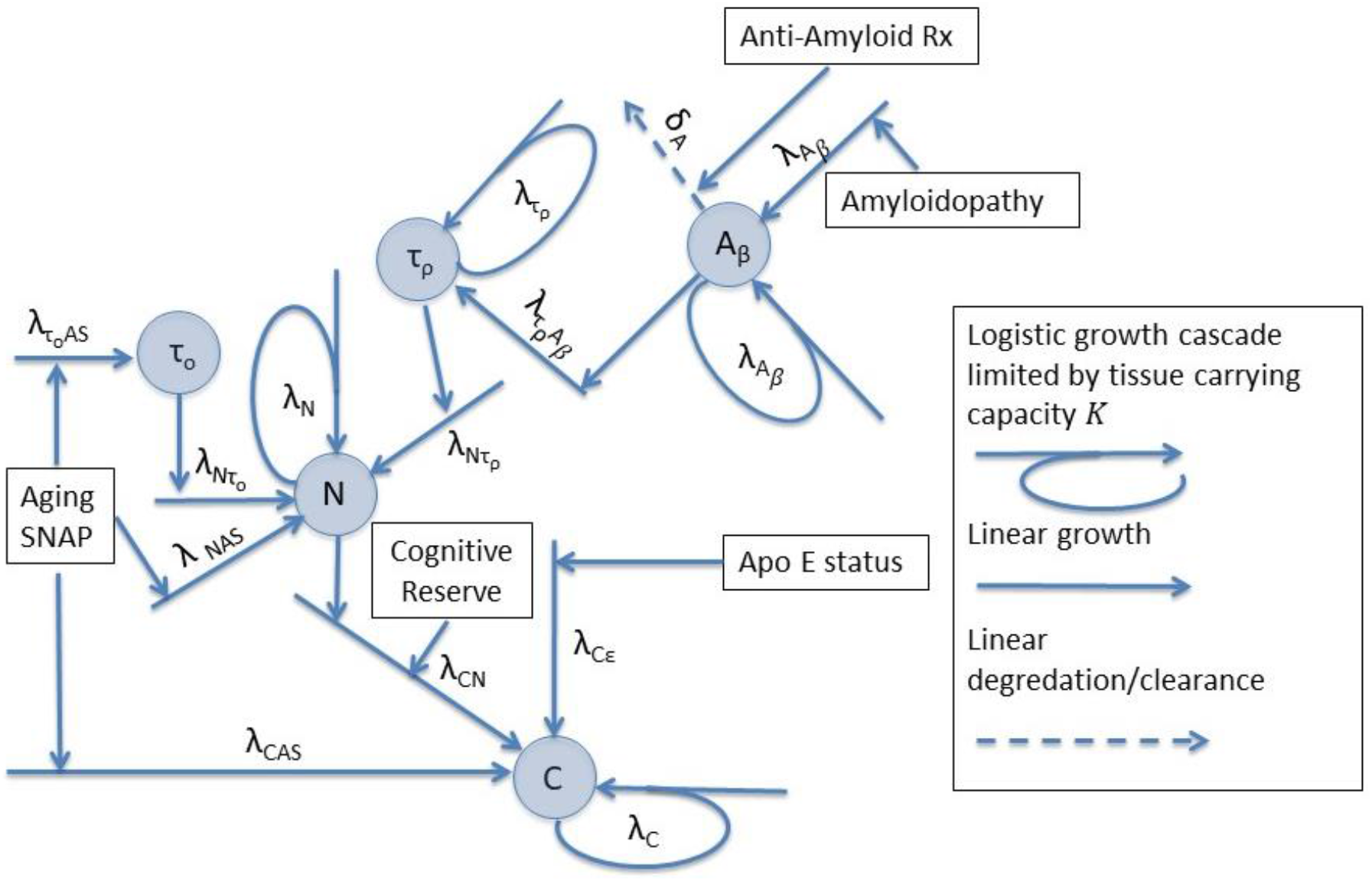
Diagram depicting causal modeling implementation of the biomarker cascade model in Alzheimer’s disease (1, 2). Blue circles represent biomarker quantities. A_*β*_ represents amyloid pathology. Its initial value determines when during the lifespan the amyloid cascade begins. τ_*ρ*_ represents amyloid-related tau pathology (p-tau). τ_*0*_ represents age-related and/or suspected non-Alzheimer pathology (SNAP)-related tauopathy. *N* represents neuronal dysfunction/loss. *C* represents cognitive impairment. λ values are growth rate constants, and δ a degredation/clearance rate constant. Amyloidopathy, aging, SNAP (suspected non-Alzheimer’s pathology), ApoE status and cognitive reserve are constants that modify the onset of the growth cascades. Anti-amyloid therapy is a function of time. Descriptions of all variables and parameters are listed in Table 1 and Table 2, respectively.© 2017 - 2018 Duke University. All Rights Reserved.

### Computational Model Construction

We implemented the above causal model, using the ordinary differential equation (ODE) toolbox in MATLAB (Mathworks^®^, Natwick, MA), as system of non-linear first order ODEs to include amyloid dependent and non-dependent neurodegenerative cascades. The amyloid dependent cascade is initiated by amyloid beta, A_*β*_, and mediated via phosphorylated tau, τ*_ρ_*. The non-amyloid-dependent cascades are initiated by comorbidities, e.g., aging and/or suspected non-Alzheimer pathology (SNAP), either directly, or indirectly via non-amyloid dependent tauopathy, τ*_0_*,. Initiation of cognitive decline, *C*, is directly determined by neurodegeneration, comorbidities, genetic factors and cognitive reserve. The equations are as follows (© 2017 - 2018 Duke University. All Rights Reserved):

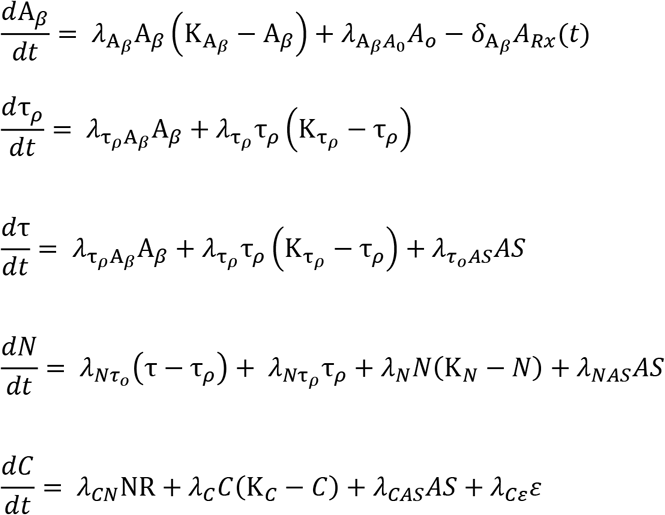

where A_*β*_ represents amyloid pathology, τ*_ρ_* represents amyloid-related tau pathology (p-tau), τ_*0*_ represents age-related and/or SNAP-related tauopathy, τ represents total tau pathology, defined as the sum of τ*_ρ_* and τ_*0*_, *N* represents neuronal dysfunction/loss, and *C* represents cognitive impairment. λ defines numerous rate constants λ_A_*_β_*, λ_τ_*_ρ_*, λ*_N_* and λ*_c_* reflect the logistic growth rates of the various biomarker cascades. The remaining rate constants reflect linear growth rates of the biomarkers and determine the influence of various factors on the time-of-onset of the subsequent biomarker cascades. λ*_CN_*, for example, is a rate constant that reflects the influence of neurodegeneration, which is modified by cognitive reserve, for example education level. This, along with comorbid pathologies and genetic risk alleles, determines the age of onset of the cognitive decline cascade. *δ*_A_*_β_* represents a degredation rate constant for A_*β*_, and in this model, mediates the effects of anti-amyloid therapy. A*_0_* represents amyloidopathy, *A_Rx_* (t) represents a time-dependent function for anti-amyloid therapy, AS represents aging and/or SNAP, R represents cognitive reserve, and ε represents ApoE allele status. Descriptions of all variables and parameters are listed in Table 1 and Table 2, respectively.

**Table 1:** Model Variables

| Variable | Description |
| --- | --- |
| $A_{\beta}$ | amyloid beta pathology |
| $\tau_p$ | amyloid-related phospho tau pathology |
| $\tau_o$ | SNAP-related tau pathology |
| $N$ | neuronal degeneration |
| $C$ | cognitive impairment |

SNAP = suspected non-Alzheimer's pathology

**Table 2:**
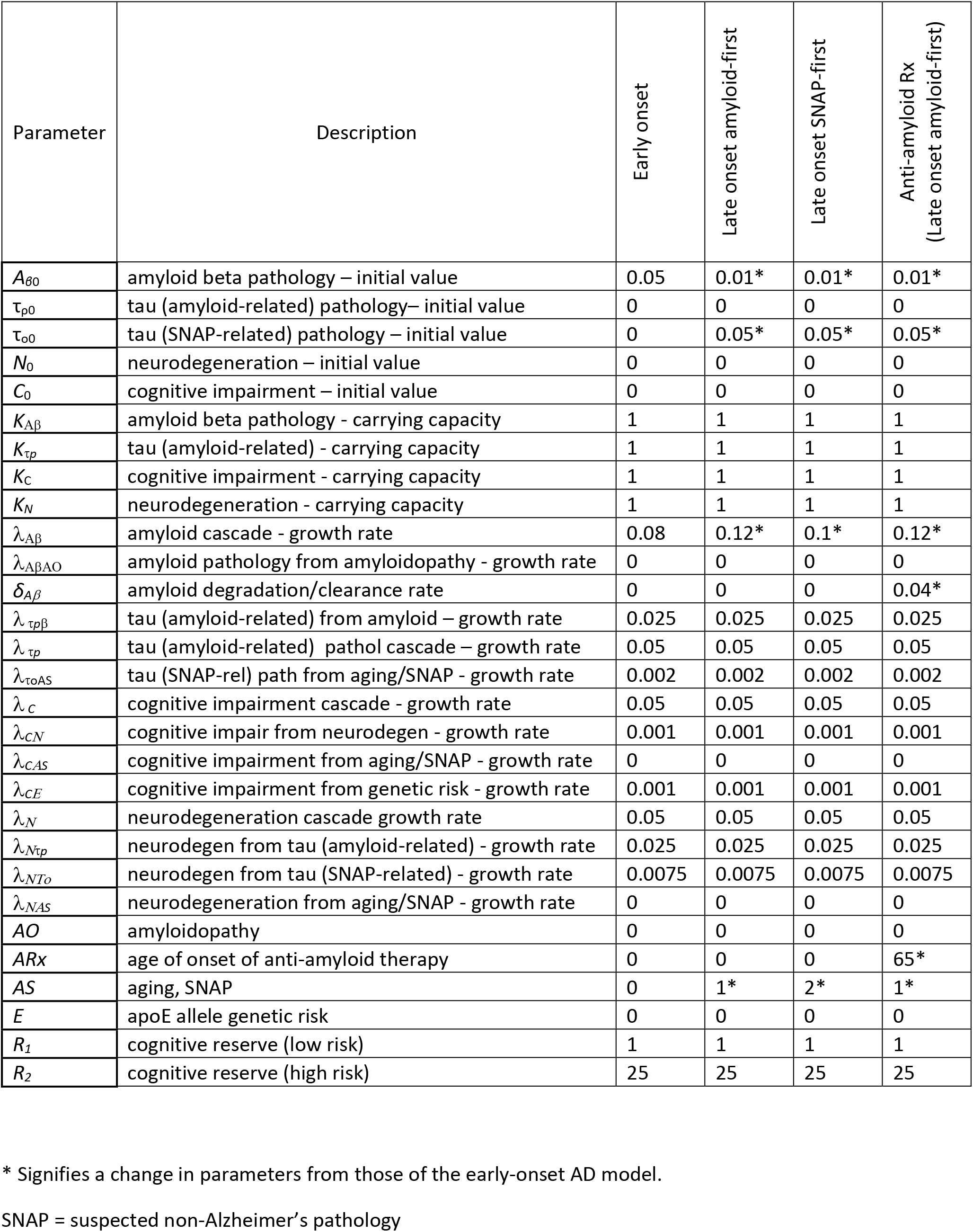
Model parameters for early- and late-onset AD and anti-amyloid therapy scenarios

Additional assumptions of the model are as follows: 1) Biomarker cascade growth is implemented via a logistic growth model with carrying capacity K. K is adjusted using a least squares minimization procedure so all biomarkers achieve maximal level of 1 at age 100 years. This is done to configure biomarker curves in a sigmoidal shape with a progressively steeper slope in the right-hand tail for later changing biomarkers, as described in the hypothetical model; 2) At time t=0, A_*β*_ is set to a very small number. This is done to initiate the amyloid cascade sometime during the lifespan, even in the absence of amyloidopathy. For simplicity, amyloidopathy A*_0_*, is set to zero in all models, and a slightly larger initial value of A_*β*_ is used for the early-onset model, whereas a slightly smaller value is used in the late-onset models. All other biomarker initial values are set to zero, except for total tau in the late onset models, which is set to the minimum biomarker level on the graphs; 3) The minimal biomarker level on the graphs is set to 0.1 to allow for different onset-delays for the sigmoidal shaped biomarker curves that depend upon both biology and biomarker sensitivity; 4) Amyloidopathy, SNAP, aging, and ApoE status are constants across the age span that add linearly to the growth rate of the biomarkers and cause earlier initiation of the amyloid, tau, neurodegenerative and/or cognitive decline cascades; 5) Cognitive reserve is a constant that modifies the effect of neuronal degeneration on the onset of cognitive decline. A lower value is used in the low risk group, and a higher value in the high-risk group. 6) Anti-amyloid therapy, once initiated, is assumed to be maintained throughout the lifespan, and *A_Rx_* (t) is simulated as a Heaviside step function, H(n), using the half maximum convention,

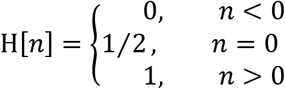

where n represents the age of initiation of therapy. In the case of anti-amyloid therapy, carrying capacity, K, was determined for all biomarkers in the no-therapy condition, in a natural history context, with *δ*_A_*_β_* set to zero. *δ*_A_*_β_* was then changed to positive number to simulate the effects of amyloid degradation, with fixed K values based on natural history. This was done to assure that the evolution of biomarkers in the pre-therapy interval was in no way influenced by the administration of therapy later in the course of the disease.

To determine the feasibility of the CCM, we parameterized and tested four versions: 1) familial autosomal dominant AD, 2) late-onset amyloid-first AD, 3) late onset tau-first AD, and 4) anti-amyloid therapy in late-onset amyloid-first AD dementia. In the first three scenarios, the goal was to determine whether manipulating the CCM parameters in a physiological meaningful manner could reproduce biomarker trajectories that closely matched those visually depicted in the literature. In the fourth scenario, we determined whether the model would predict the outcome of recently failed clinical trials of anti-amyloid therapy administered in symptomatic late-onset AD (8).

## Results

### Computational model of familial autosomal dominant AD

Figure 2 illustrates that the cascade of early-onset familial AD derived from prior literature (1) (left) matches closely to the output generated by our DCM (right). Specifically, they both demonstrate the initial appearance of amyloid, followed by tau and neurodegeneration, then followed by the onset of cognitive decline. It also shows how cognitive reserve could modify the cascade. Although tau in these figures represents total-tau, in this scenario it is dominated by p-tau (amyloid-related tau). Model parameters are shown in Table 2.

**Figure 2:**
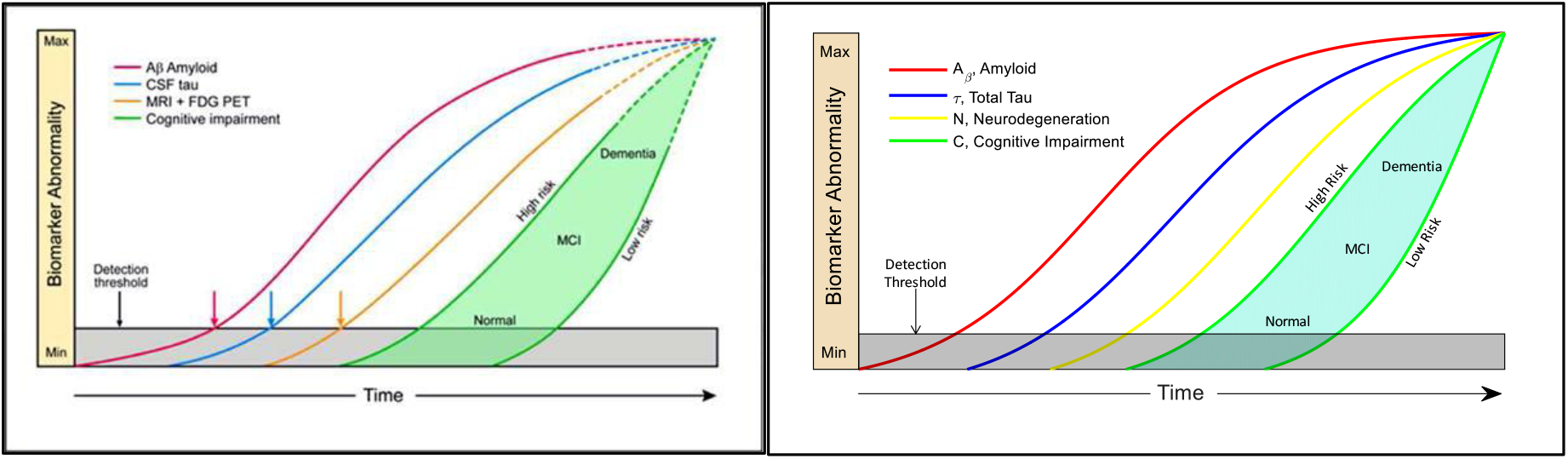
Model of early-onset, autosomal dominant AD. The red, blue and yellow curves represent the evolution of amyloid, tau and neuronal biomarker levels, respectively, over the course of the disease. The green curves represent cognition in two hypothetical high and low risk groups based on low and high cognitive reserve. Our CCM-generated curves (right) closely match the schematic model curves on the left (adapted from (1) with permission) from prior literature.

### Computational Model of beta-amyloid-first late-onset AD

The late-onset AD CCM (Figure 3) output shows that amyloid appears first, followed by total tau and neurodegeneration. In this CCM, the arrival of amyloid is delayed compared to that in early-onset AD but reaches detection threshold prior to total tau. The CCM trajectories visually matched those predicted in the literature (1). Model parameters are shown in Table 2.

**Figure 3:**
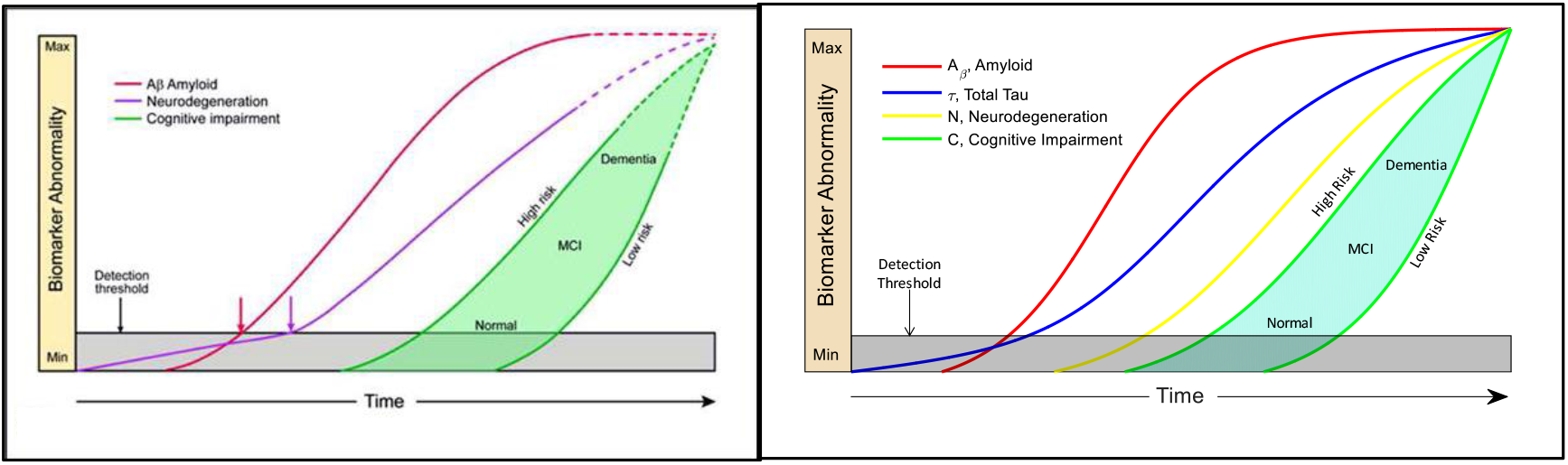
CCM of late-onset, amyloid-first AD (right panel) matches closely the pattern hypothesized in the literature (left panel - adapted from (1) with permission). The red, blue and yellow curves represent the evolution of amyloid, tau and neuronal biomarker levels, respectively, over the course of the disease. In the left panel, the blue and yellow lines are combined into a single purple line, per the original theory in which tau was considered a neurodegenerative marker. The green curves represent cognition in two hypothetical high and low risk groups, based on cognitive reserve

### Computational model of delayed beta-amyloid late-onset AD

In this CCM (Figure 4), the arrival of total-tau precedes that of amyloid and initiates neurodegeneration, whereas the subsequent appearance of amyloid accelerates this process. Our CCM mimics a condition described in literature as suspected non-amyloid pathology (SNAP) in some ways (absence of initial amyloid) but illustrates a mixed pathology concept, where amyloid and amyloid-related tau contribute to cognitive decline at later stages. Model parameters are shown in Table 2.

**Figure 4:**
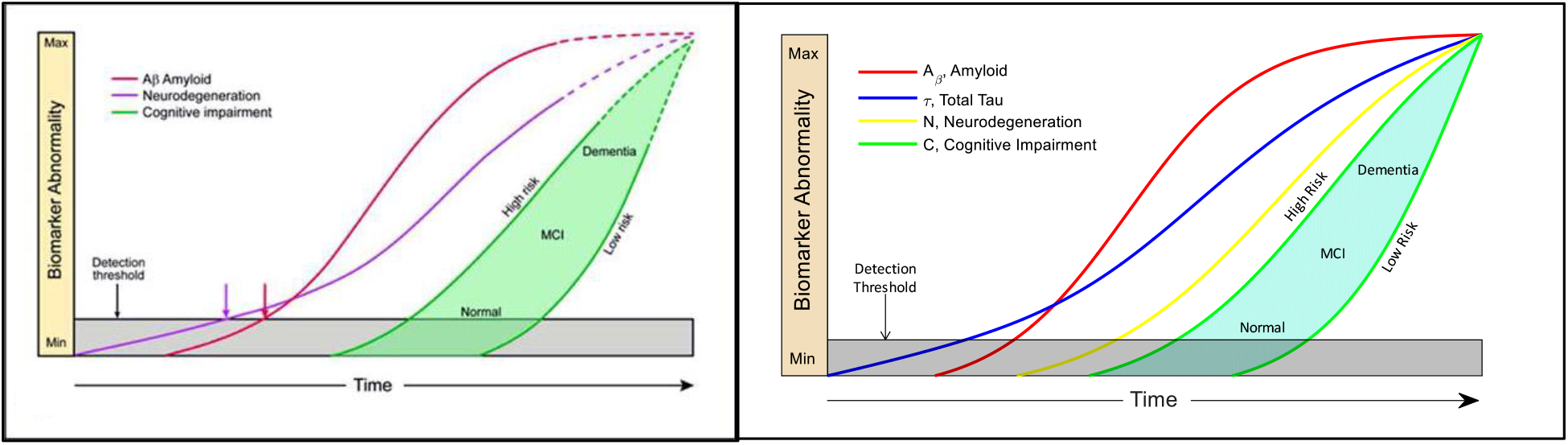
The red, blue and yellow curves represent the evolution of amyloid, tau and neuronal biomarker levels, respectively, over the course of the disease. In the left panel (adapted from (1) with permission), the blue and yellow lines are combined into a single purple line, per the original theory in which tau was considered a neurodegenerative marker. The green curves represent cognition in two hypothetical high and low risk groups, based on cognitive reserve. The CCM (right panel) closely matches the trajectories proposed in the literature.

### Simulation of anti-amyloid therapy administered in amyloid-first late-onset AD dementia

Figure 5 depicts the outcome of our CCM of anti-amyloid therapy when given to amyloid-first late-onset AD dementia patients after symptom onset. This model output shows no benefit on the onset or slope of cognitive decline, despite the amyloid level dropping substantially from its peak. Tau levels drop marginally. The model mimics the results of recent failed anti-amyloid therapy trials in probable AD dementia (8). In this model, anti-amyloid therapy would have to be given before a hypothetical tipping point to show benefits on cognition. Model parameters are shown in Table 2.

**Figure 5:**
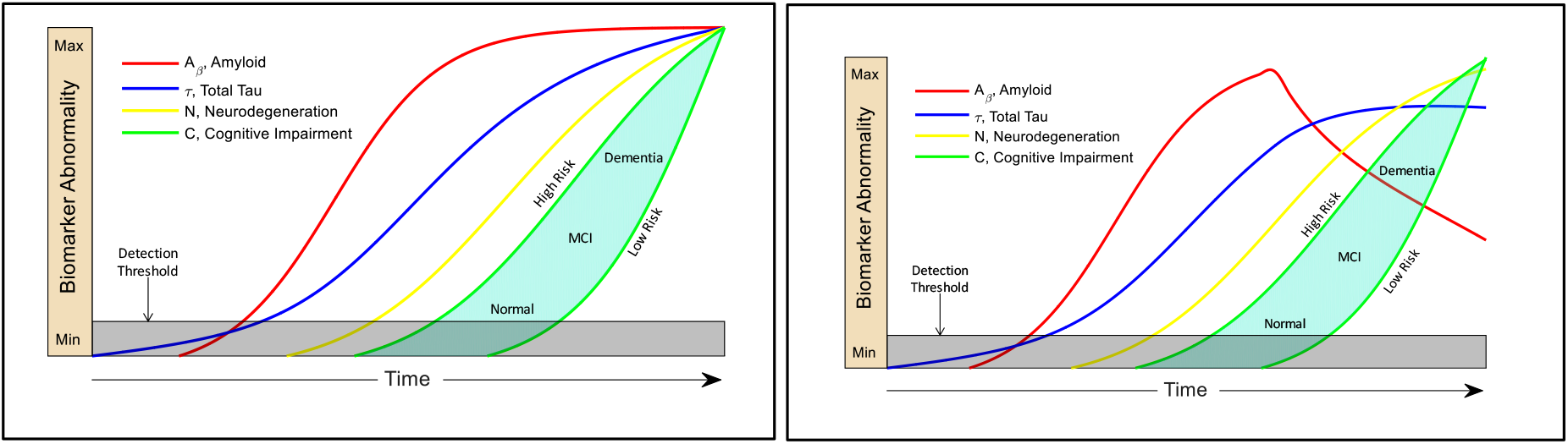
Left panel shows the untreated condition for comparison, reproduced from Figure 3. Right panel showing the CCM simulation of the effect of anti-amyloid therapy administered in AD after symptom onset. The red curve shows marked decline in brain amyloid levels, the blue line shows a small decrease in tau levels and green lines show there is no significant effect on cognitive decline onset or rate, consistent with the many failed trials.

## Discussion

We have implemented a CCM that incorporates the three clinically available categories of biomarkers to track AD progression, amyloidopathy, tauopathy, and neurodegeneration. The model effectively simulates the temporal evolution of the biomarkers and their relation to cognitive decline as described in previous literature (1, 2, 5), taking into account late verses early onset, the influence of aging and cooccurring non-AD-related brain pathology common in the elderly, and the concept of cognitive resilience to AD pathologic changes. In addition, we simulate the effects of a disease-modifying therapy given late in the disease course, after patients become symptomatic. This CCM was developed both as a means to test existing theories and as a new resource for the field that can be refined as our knowledge advances.

The hypothetical model of the AD pathological cascade, originally published in 2010 (5), and updated in 2013 (2), is based largely on cross-sectional biomarker data, due to limited individual longitudinal biomarker data. It postulates a temporal evolution markers of amyloid pathology, tau pathology, and neurodegeneration, represented as sequential plots of biomarker abnormality over time, leading to cognitive impairment. Three different pathological and neuronal loss scenarios were considered, early onset familial AD, amyloid-first late onset AD, and non-amyloid first (SNAP) late onset AD (1). We created and parameterized a CCM, based on assumptions of underlying biology inherent in the AD pathological cascade, to successfully simulate these three natural history scenarios. Our CCM of early-onset autosomal dominant AD closely matched the temporal order and shape of the biomarker trajectories in the literature schematic (5), and is also supported by empirical data from the longitudinal Dominantly Inherited Alzheimer Network (DIAN) study (9). Our CCM of amyloid-first late onset AD and SNAP-first model of AD also closely simulated the curves postulated in the literature (2). Lastly, our simulation of anti-amyloid therapy in late-onset AD dementia mimicked the negative findings from several failed clinical trials of anti-amyloid therapies (8). Of note, most of the disease modifying treatment trials in preclinical disease or in mild AD, both several recently completed and ongoing, target beta-amyloid or tau pathologies.

There are some limitations to our work. First, the hypothetical model (1) on which we built our CCM may not be entirely accurate. Although, some aspects AD pathological cascade model, for example the ordering and shape of the biomarker curves, have been validated using data-driven approaches (10, 11), the model continues to evolve as more natural history and clinical trial data becomes available. Second, we did not parameterize our models using actual patient biomarker data. Rather, our goal in this work was to construct and test the feasibility of a CCM that would best match the hypothetical biomarker trajectories proposed in the literature (1). Subsequent goals will include adjusting parameters based on longitudinal data and iteratively refining the model itself as new knowledge becomes available. Third, our CCM is a simplified causal model of biomarkers interacting with each other, an abstraction that does not model the actual underlying cellular and molecular processes. Prior CCM efforts in AD have modeled the disease at a molecular and cellular level (12, 13) as well as at a whole-brain, systems level using MRI and EEG data (14, 15). There have been few CCM approaches that have specifically focused on clinical AD biomarkers (14, 16), and none have incorporated all three clinically available categories of biomarkers, amyloidopathy, tauopathy, and neurodegeneration. Overcoming the above limitations will require large real-world data sets of individual longitudinal biomarker trajectories across the cognitive continuum as well as integration of genomic, cellular and biomarker knowledge. Such efforts are underway on an international scale (17–20).

A key strength of the CCM is that it allows for testing underlying causal assumptions in an integrated fashion, unlike other published correlative mathematical models of clinical biomarkers (10, 11, 21–27) that treat them independently or fit the data without considering its underlying causal structure. For example, several studies validated the temporal ordering of biomarkers, without attempting to explain the underlying disease mechanism by which this temporal ordering arises (10, 11, 26, 27). Causal models allow for testing the effects of non-linear interactions among multiple AD biomarkers and comorbid conditions that cannot be deduced by intuition alone, as well as for predicting response to single and combination therapies. A CCM can be implemented in a “forward” manner to simulate new data or in a “backward” manner, using a Bayesian inversion procedure, to infer the causal architecture of the system based on existing data. This approach has been applied extensively to re-construct mechanistic models of brain function and disease, including AD, from electrophysiologic and imaging data (14, 16, 28). Unlike descriptive models of disease, which can be become increasingly difficult to validate, particularly as datasets of biomarker trajectories become larger and more complex, CCM’s can be easily scaled up to increasing degrees of complexity. It is our hope that once a CCM resource for clinical AD biomarkers is created, it will be parameterized based on data from patient studies, expanded and iteratively refined over time. Ultimately such a model would create a global resource for the field to translate existing knowledge, personalize care and accelerate drug discovery for this devastating disorder (29).

